# Habitat heterogeneity reduces richness of ant species by increasing abundance of the local dominant species

**DOI:** 10.1101/316430

**Authors:** B. Travassos-Britto, P. B. L. Rocha

**Author notes:** Corresponding author: Address: Rua do Matão, 321 - Travessa 14, Cidade Universitária - São Paulo - SP, Brazil, 05508-090.

## Abstract

The effect of environmental heterogeneity on species richness is frequently discussed in ecology. However, the empirical evidence has been contradictory as to the direction of the effect. Although some authors have considered that this might be a methodological problem, we argue that for ants, ecological interactions within the community, as interspecific competition is more important. We analyzed the plausibility of models in explaining the ant richness distribution patterns in a semi-desert environment. We used three predicting variables in the construction of the models to explain ant richness distribution: heterogeneity based on the amount of structures regardless of their type, heterogeneity based on the diversity of structures, and the abundance of individuals of the dominant species. We used ANOVA to chose the best model and corroborated the prediction that in this system abundance of dominant species is the best predictor of ant species richness. Neither of the heterogeneity conceptions contributed much to explain richness distribution. However, in a second analysis, we concluded that heterogeneity could affect the abundance of the dominant species. We conclude that competitive dominance is a better predictor of species richness distribution patterns than structural heterogeneity. However, the structural heterogeneity affects the distribution of dominant individuals. We suggest that some unexplained patterns observed about the relationship between heterogeneity and richness could be due to an indirect effect.

## Resumo

O efeito da heterogeneidade ambiental na riqueza de espécies é frequentemente discutido na ecologia. Entretanto, as evidências empíricas têm sido contraditórias com relação à direção do efeito. A pesar de alguns autores considerarem que essa divergência é causada por motivos metodológicos, nós argumentamos que para formigas, interações dentro da comunidade, como competição interespecífica é mais importante. Nós analisamos a plausibilidade de diferentes models em explicar o padrão de distribuição de riqueza de morphospecies de formigas em um ambiente semi-desértico (caatinga). Nós usamos três variáveis preditivas na construção dos modelos para explicar a distribuição de riqueza das espécies: heterogeneidade baseada na quantidade de estruturas independentemente dos tipos, heterogeneidade baseado na diversidade de estruturas e abundância de indivíduos da espécies dominante de formiga. Nós usamos ANOVA para escolher o melhor modelo e corroboramos a predição que nesse sistema a abundância da espécie dominante é o melhor preditor de riqueza de espécies de formiga. Nenhuma das concepções de heteorgeneidade pareceu contribuir muito para explicar a distribuição de riquezas de formigas. Entretanto, em uma segunda análise, concluímos que heterogeneidade tem efeito sobre a abundância da formiga dominante. Nós concluímos que a dominancia competitiva é um melhor preditor do padrão de distribuição de riqueza das espécies que a heterogeneidade de estruturas. Entretanto, que heterogeneidade de estruturas influencia a distribuição de indivíduos dominantes. Sugerimos que alguns padrões não explicados observados sobre a relação entre heterogeneidade e riqueza pode ser explicado por efeito indireto sobre padrões mais importantes.

The distribution patterns of species richness can be influenced by external structures, which affect niche availability in an environment (MacArthur & MacArthur, 1961; Tews et al, 2004), an important property of the environment. The distribution of species within a community can also be affected by how interspecific interactions occur among species (Andersen, 1992; MacArthur & Levins, 1964; Parr, 2008; Savolainen & Vepsäläinen, 1988), a property of the community itself. The concept that changes in a habitat’s physical structure can cause changes in the distribution of species in space is not new in the context of ecological studies. The extensively explored niche theory states that the distribution of species is driven by the interaction of individuals with the characteristics of their environment (Hutchinson, 1959). The effects of the environment on organismal distribution can be deconstructed into the following two types: the effect of the physical structure of the environment and the effect of interspecific interactions (Soberón, 2007). There are many models that explain variation in species distributions based on structural variation and many others that explore the effect of interspecific interactions. These two types of models lead to explanations as to why some sites have more species than others that are based on different mechanisms. Here, we compare the plausibility of models based on both mechanisms to explain the distribution of ant richness in a semi-desert environment.

The habitat heterogeneity hypothesis suggests a mechanism in which structural variation affects species distribution (Pianka, 1966), and it is a well-discussed theme in ecology (Heck Jr & Wetstone, 1977; MacArthur, 1964; Pianka, 1966; Simpson, 1964; Tews et al, 2004). The hypothesis predicts that more heterogeneous habitats will have higher species richness (MacArthur & MacArthur, 1961). This hypothesis is based on the argument that heterogeneous habitats have more diverse structural arrangements and therefore can be exploited in more ways, allowing the coexistence of a greater number of species via a reduction in competitive pressure. However, this model is not completely supported by empirical data (August, 1983; Cramer & Willig, 2002; Hill et al, 1995; Kotze & Samways, 1999; Lassau & Hochuli, 2004; Lassau et al, 2005; Tews et al, 2004; Wiens, 1974). Because the mechanism underlying the heterogeneity hypothesis is very intuitive, researchers have found it difficult to understand cases in which the heterogeneity hypothesis is not supported by empirical data and to determine what mechanisms are acting in these cases (Tamme et al, 2010; Tews et al, 2004; Travassos-De-Britto & Rocha, 2013).

Different authors have suggested that such controversial findings may result from problems with the methodologies of the studies that have tested the heterogeneity hypothesis. One of the most notorious methodological problems with testing the heterogeneity hypothesis is how heterogeneity itself is determined; different measures of structural heterogeneity may lead to contradictory results concerning its association with species richness (Heck Jr & Wetstone, 1977; Seibold et al, 2016; Travassos-De-Britto & Rocha, 2013). Therefore, a proper assessment of the effect of heterogeneity on species distribution patterns should take this issue into consideration.

The two most common concepts of structural heterogeneity proposed in recent studies are heterogeneity defined either as the amount of sensible structures regardless of their characteristics (dubbed hereafter as Heterogeneity 1) or as the diversity of structural elements, taking into consideration how many types of structures there are in the environment (Heterogeneity 2) (see Tews et al, 2004, for a review). The mechanism explaining the positive relationship between heterogeneity and richness is slightly different depending on the type of heterogeneity considered. For Heterogeneity 1, the greater the amount of structures, the higher the probability of generating more structurally diverse microhabitats, allowing for different species to make use of these microhabitats. In a more homogeneous environment (i.e., with less microhabitats), competitive pressure is higher, leading to fewer species. For Heterogeneity 2, the diversity of structures directly changes the number of ways it is possible to exploit the environment. A greater diversity of structures could mean a greater diversity of shelter, food, and nesting places (see Travassos-De-Britto & Rocha, 2013). Because models based on different concepts of heterogeneity might be explained by different mechanisms, we should treat them as two different predictive variables.

Alternatively, the contradictory results concerning tests of the heterogeneity hypothesis might be due to non-methodological issues. Other variables not associated with the structure of the environment could be driving species distribution patterns. It is difficult to determine what could be more important than heterogeneity to species richness in every system. However, in some systems, there are variables that are especially important. Studying these systems could be a convenient way of assessing the importance of heterogeneity in determining species distribution patterns.

Competition is an interspecific interaction and has being studied for a long time (MacArthur & Levins, 1964). Although some studies have shown that competition may not be as important in shaping communities as previously thought (Connell, 1980; Hubbell, 2001), in some systems, competition continues to be considered of great importance in understanding species distribution patterns (Cerda et al, 2013; Ligon et al, 2011; Sanders et al, 2003; Vahl et al, 2005; van Klink et al, 2015). For example, ant assemblages have a structure based strongly on competition (Cerda et al, 2013; Hölldobler & Wilson, 1990; Parr, 2008). Competition has been shown to shape behavioural and spatial distribution patterns in ant communities (Vahl et al, 2005; Vepsäläinen et al, 2000). In ant assemblages, there are species that are markedly more abundant than others (numerically dominant) or that exhibit a more aggressive foraging behaviour (behaviourally dominant). Arnan et al (2011) observed that numerically dominant species will often exclude other species by quickly extinguishing the resources at a site and that behaviourally dominant will exclude other species by directly attacking ants from other nests (Arnan et al, 2011; Parr, 2008; Segev & Ziv, 2012). In a specific territory, the non-dominant species are usually referred to as the submissive ant species. Submissive species will seldom be found foraging at the same sites as dominant species (Arnan et al, 2011). Searching for resources within the dominant species territory is energetically risky. Either because they are numerically superior and therefore have a much higher probability of finding and consuming food or because they are more aggressive and will kill stray foragers from other nests. Therefore, the submissive species exploit the site’s resources by avoiding the dominant species (e.g., by quickly consuming incoming resources before the dominant species arrives at the site, by foraging at a different time of the day or by avoiding the chemical trails of dominant individuals) (Hölldobler & Wilson, 1990). This type of dynamic leads to a model that predicts species distribution patterns based on the presence of the dominant species Parr (see 2008); Savolainen & Vepsäläinen (see 1988, for empirical evidence).

The importance of competitive dominance in ants may indicate that for this taxon, the dominant-submissive dynamics in the community might be more important than habitat heterogeneity. Although there has been evidence of the positive effect of heterogeneity on ant species richness (e. g. Bestelmeyer & Wiens, 2001; Perfecto & Snelling, 1995), other studies have also shown no effect or even negative effects (Feller & Mathis, 1997; Lassau & Hochuli, 2004).

There are no reasons to think that the mechanism used to explain how heterogeneity affects species richness does not explain ant richness distribution patterns. Different species of ants should have a minimum degree of differentiation in resource necessities, and more heterogeneity should favour coexistence, thereby increasing local species richness. However, this type of dominance relationship structure occurs frequently in ant assemblages worldwide (Hölldobler &Wilson, 1990), indicating that this characteristic is strongly linked to the Formicidae family and that it likely has a strong influence on species distributions within an ant assemblage. Despite these divergent expectations about the effect of heterogeneity and presence of dominant on richness distribution, there are no studies that statistically compare the contribution of these variables in models to explanation ant richness distribution.

Here we developed different models explaining distribution of richness by Heterogeneity 1, Heterogeneity 2, abundance of dominant species or a combination of these variables. The objective of this study was to determine which model best explains the distribution of ant species in the dunes of northeast semi-desert environment Brazil.

The dunes of this semi-desert environment can be considered an extreme environment for ants. Most of the environment is exposed sand that reaches extremely high temperatures during day, with sparse patches of vegetation. These characteristics could accentuate the effect of structural heterogeneity because small changes in structure could produce sharp changes in microhabitats. However, as a semi-desert environment, it also provides few resources, which should accentuate competition (Cramer & Willig, 2002). The amount of structures should be especially important in this extreme environment. For example, a larger patch of leaf-litter can hold more humidity than a small one. However, a larger patch also requires more energy to walk through if the heat is not a problem (e. g. for night-time species). The diversity of structures should also be important because the types of structures present are very different from one another (e.g., cacti, arboreal and shrub plants, bromeliads, exposed sand, leaf-litter). Because these are characteristics that should accentuate the effects of both types of heterogeneity, we are including these two variables in the models building. However, we expected that the dominance model would better explain the richness of ant species because competition has been demonstrated to be a very important driver of ant richness distribution patterns in many communities and environments.

## METHODS

### Study Area

We conducted the study in a sand dune region in northwest Bahia, Brazil. These dunes are located along the middle of the São Francisco River valley. The climate of this region is described as arid to semi-arid (Barreto, 1996). High temperatures, with an annual mean air temperature exceeding 26.2°C and soil temperature exceeding 50°C during the day, make this place an extreme environment. This region is included in the caatinga morphoclimatic domain (bahia, 1978). The vegetation physiognomy presents trees and bushes that are short and scattered and lacks conspicuous herbaceous cover, even in the wet season. Therefore, most of the sandy soil remains exposed, except for patches of the terrestrial bromeliad *Bromelia antiacantha* (Bertol.), the small cactus *Tacinga inamoena* (K. Schum.) N.P. Taylor and Stuppy, and ground litter. We selected this area because its structural habitat elements can be easily measured, and the harsh environment of the dunes should enhance the effects of structural heterogeneity on microhabitat variables.

### Ant sampling

Ant sampling was carried out on three different days of a year. The first day of sampling was at the peak of the wet season (February), the second was at the peak of the dry season (September) of the same year, and the third was at the peak of the wet season of the next year. We distributed 119 pitfall traps on each day of sampling. These pitfall traps were arranged in a sampling grid with 17 lines and 7 columns placed 10 m apart, with a total area of 11,200 m^2^ per grid. The grids for each day of sampling were plotted at a distance of least 250 m from where the grids from the previous sampling days were plotted. Each pitfall trap consisted of three radial 1.5 × 0.4-m plastic drift fences converging on a 20-L dry bucket. Ants were removed from the pitfalls at dawn and at dusk and were immediately preserved in 70% ethanol and brought to the laboratory for screening and morphospecies identification. Pik et al (1999) has demonstrated that the identification of morphospecies closely reflects the species identification of ants.

Because ants were collected in a grid spatial autocorrelation could mask the effect of the variables of interest in our study. We tested for the effect of spatial autocorrelation on richness among sampling grids using with Moran’s I (Diniz-Filho et al, 2003).

### Measuring variables

In the study area, the following six easily identifiable types of structures were used to assess heterogeneity: leaf litter, a terrestrial bromeliad species (*Bromelia antiacantha*), a small cactus species (*Tacinga inamoena*), shrubs, subshrubs and trees. Each of these types of structures are potentially different structures to ants (Lassau et al, 2005; Rocha et al, 2010). The bromeliads can accumulate water in ponds in their centre, which may attract other arthropods. The cacti produce flowers and succulent fruit but produce very little shade. Sub-shrubs alone do not offer protection from the sun but may serve as food and shelter from some predators. Shrubs provide areas of higher humidity and protection from the heat and against terrestrial predators such as lizards or rodents. Trees can offer more nesting sites and considerably more leaves but seldom provide protection against visually oriented predators. Leaf-litter may offer protection and increased humidity but is also more difficult to move across than bare sand (Hughes & Ward, 1993).

To measure habitat heterogeneity in each sampling unit, we first drew a 3-m-diameter circle centred in each pitfall trap. Then, we measured the projected area of each of the six types of structures in mm^2^. The areas of shrubs and trees were computed by summing the projected area of each individual plant, considering overlapping of projection. For a graphical depiction of the structural measurements, see Figure 3 in Rocha & Rodrigues (2005).

We used two different indexes of heterogeneity: Heterogeneity 1 and Heterogeneity 2. Heterogeneity 1 was determined by summing the coverage area (mm^2^) of the structures in each trap unit, and Heterogeneity 2 was determined based on the diversity of structures in each trap unit, as defined by the Shannon-Weiner diversity index. The spatial scale adopted was intended to allow for a considerable amount of heterogeneity among the sampling plots that could be perceived among the populations of ants. The habitat attributes chosen to reflect heterogeneity were those related to plant growth patterns, which have a close relationship with ant ecology (Beattie, 1985; Brener & Silva, 1995; Hölldobler &Wilson, 1990; Leal & da Silva, 2003).

We used the abundance of the dominant morphospecies as and index of dominant species presence. We identified the dominant morphospecies based on differences in abundance and occurrence, i.e., the species that occurred disproportionately more than others (Cerda et al, 2013; Hölldobler &Wilson, 1990; Segev & Ziv, 2012).

#### Statistical Analysis

We compared models that predicted ant richness distribution based on combinations of the three variables Heterogeneity 1 (amount of structures), Heterogeneity 2 (diversity of structures) and abundance of the dominant species. Because we think the day of sampling might be a confusion factor and we wanted to isolated the effect of this variable we included the day of sampling as the random effect in all models. Because the error distribution of our dependent variable (morphospecies count) followed a typical distribution for count data, and because we had a high number of sample units with no ants, we used Zero Inflated Poisson Generalized Linear Mixed Models (ZIPGLMM) (Zuur et al, 2009).

We also analysed whether the abundance of the dominant species was influenced by heterogeneity. This analysis had the potential to indicate the existence of an indirect effect of heterogeneity on species richness via an influence on the abundance of the dominant species. For this analysis, we generated ZIPGLMM with the abundance of dominant species as the response variable and combinations of Heterogeneity 1 and Heterogeneity 2. The day of sampling was included as random effect in all models.

In both model selections analysis we started with the most complex model and dropped terms that did not contributed significantly to the explanation of the model. The models with and without the selected term to be dropped were compared with ANOVA (significance level = 0.05). At the end of each selection the selected model was compared to a null model of the poisson distribution.

## RESULTS

Ants were captured in 196 of the 351 pitfall traps installed. We captured 999 ants and identified 18 morphospecies, which seems reasonable in comparison with the most extensive survey of ant species in caatinga phytophisiognomy (seeLeal et al, 2003). A total of 653 individuals (~ 65%) were identified as being from morphospecies 1. In addition, we observed morphospecies 1 in 148 of the 196 sample units that had ants (~ 70%). Figure 1 shows a comparison of the abundance and occurrence between morphospecies 1 and the other morphospecies. We designated morphospecies 1 as the dominant species because its abundance and occurrence were both disproportionately higher than those of the other morphospecies.

**Figure 1.**
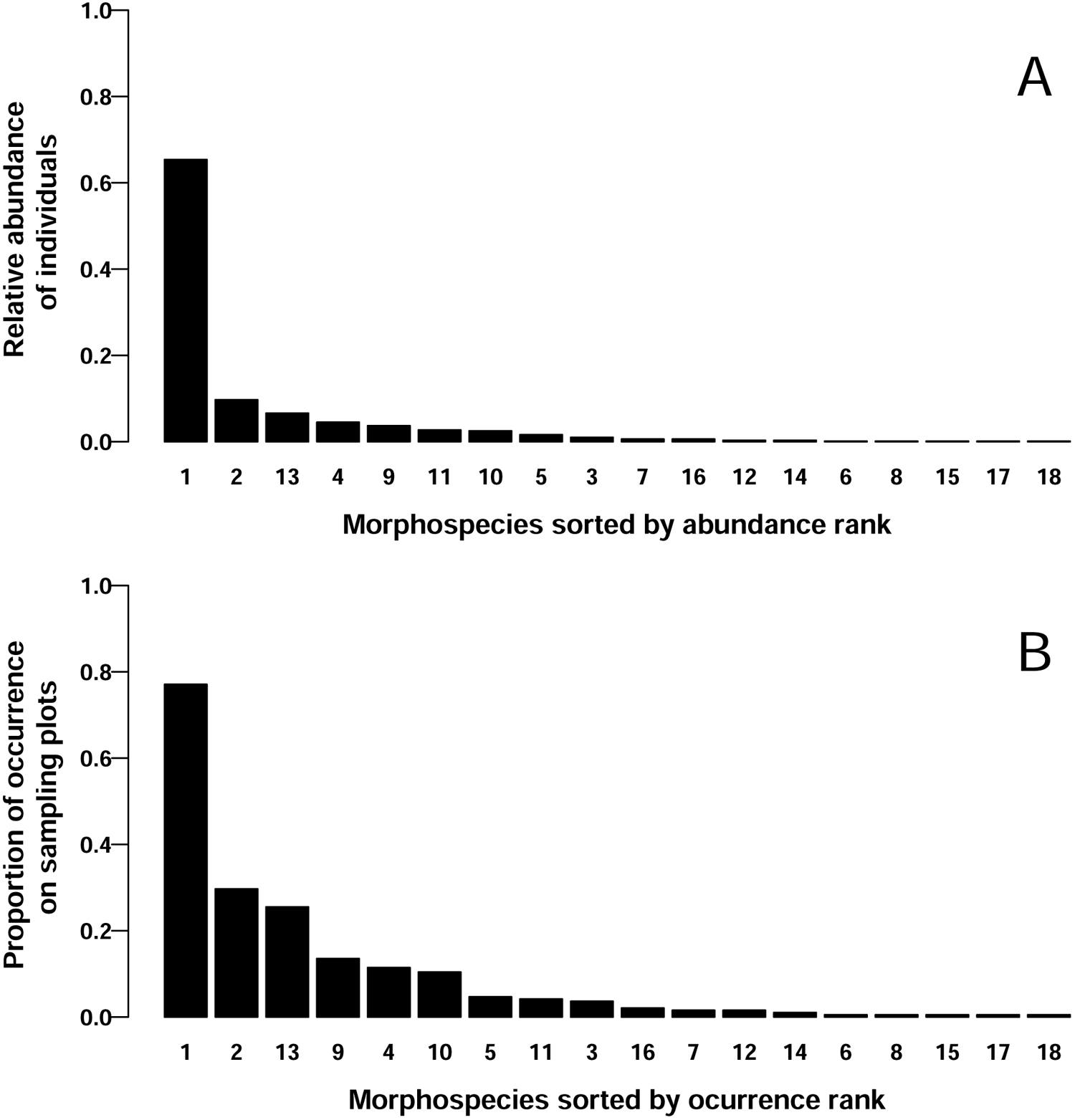
Relative abundance of individuals of each morphospecies in a total of 999 individuals (A) and the proportion of each morphospecies in the 196 pitfall traps considered (B). The numbers on the x axis indicate the identity of the morphospecies.

The Moran’s I test showed no effect of the spatial autocorrelation over richness of species in any sampling grid: Day I (Moran’s I = 0.53, *p-value* = 0.95) Day II (Moran’s I = 0.11, *p-value* = 0.34) Day III (Moran’s I = 0.21 *p-value* = 0.12).

### Selected models

Among the models comparing the effect of the heterogeneity and abundance of dominant species on ant richness distribution the model with only abundance of dominant species as predictor was selected 1. The abundance of dominant species presented significant effect on both the count of richness of other species and in the presence and absence of other ants. Poisson count model (*z-value*= *−*2.97, *p-value<* 0.003) and logistic model (*z-value*= *−*2.225, *p-value<* 0.026). The fit of this model to the data is depicted in Figure 2.

**Figure 2.**
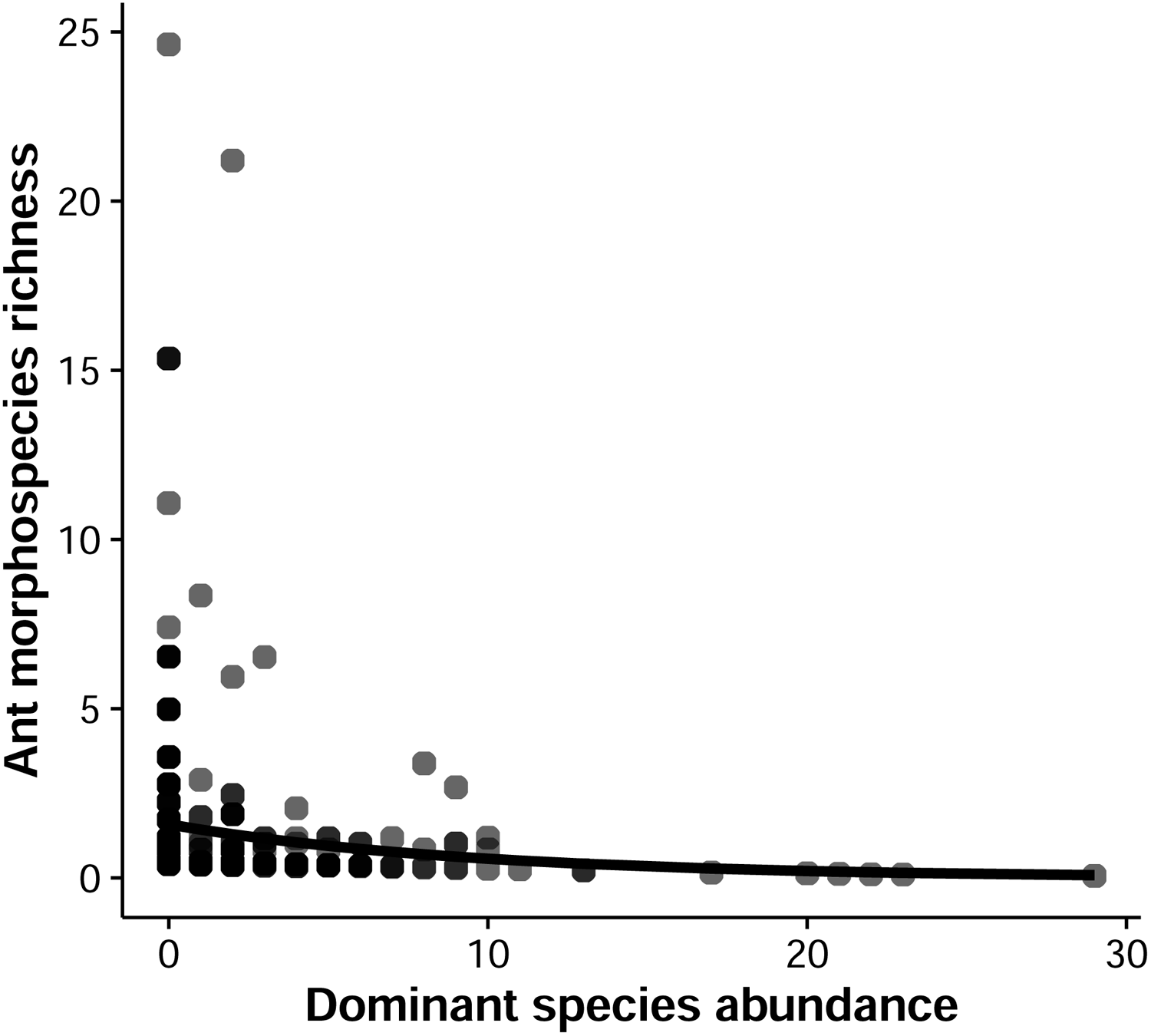
Poisson model fit to the richness of species data in relation to the abundance of dominant morphospicies. Because we wanted to present the relationship of these two variables when the effect of the day of sampling is removed, and because the link function of a poisson distribution is logarithm we used we used the exponential residuals of the model in the ‘y’ axis.

Among the models comparing the effect of the heterogeneities on the abundance of dominant species, the model in which the interaction between both measures of heterogeneity was the predicting variable was selected (Table 2). In the selected model all variables presented significant effect on the poisson count of richness: Heterogeneity 1 (*z-value*= *−*3.10, *p-value<* 0.01), Heterogeneity 2 (*z-value*= *−*5.35, *p-value<<* 0.01), Heterogeneity 1 * Heterogeneity 2 (*z-value*= 7.32, *p-value<<* 0.01). However, for the binomial model of presence and absence of dominant species only Heterogeneity 1 some effect: Heterogeneity 1 (*z-value*= 3.180, *p-value<* 0.001). The fit of the interaction model to the data is depicted in Figure 3.

## DISCUSSION

Our results indicate that the variation in richness of ants in the study site, including the sites with no ants, was explained by the abundance of the dominant species alone (see Table 1). However, heterogeneity might have an indirect effect on a richness distribution. Our results suggest that the effect of environmental heterogeneity in defining species richness distribution patterns might not be as important as previously thought when compared with the effects of interspecific interactions. Here, we discuss some possible mechanisms to explain the observed patterns.

**Table 1.** Models explaining ant richness distribution by two heterogeneity measures and dominance relationship. The models are presented in complexity order from top to bottom. The terms dropped were those with the higher p-value within each model (Zuur et al, 2009). The values in the third, forth and fifth columns are the comparison between the model in the line and the model immediately below. The selected model is highlighted in bold.

| Models | *LogLik* | *df* | *Deviance* | *p – value* |
| --- | --- | --- | --- | --- |
| Het 1 * Het 2 * Dominance | -362.75 | 2 | 1.818 | 0.40 |
| Het 1 * Dominance + Het 2 * Dominance | -363.66 | 1 | 0.052 | 0.82 |
| Het 1 + Het 2 * Dominance | -363.69 | 1 | 0.166 | 0.68 |
| Het1 + Het 2 + Dominance | -363.77 | 1 | 0.114 | 0.74 |
| Het1 + Dominance | -363.82 | 1 | 1.218 | 0.27 |
| **Dominance** | **-364.43** | **1** | **10.376** | < 0.01 |
| Null model | -369.62 |  |  |  |

**Table 2.**
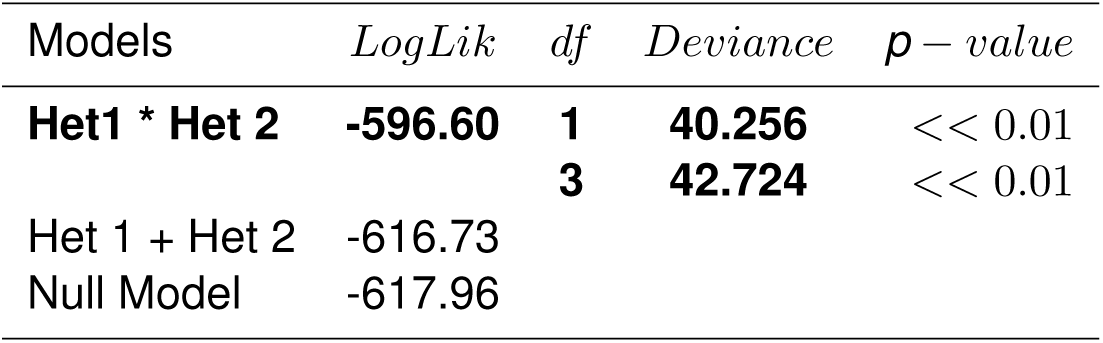
Models explaining dominant ant abundance by two different heterogeneity measures. In the first line there is the comparison between the model with interaction between heterogeneity measures and the model without interaction. The second line shows the comparison between the interaction model and a null model. The interaction model was selected.

| Models | *LogLik* | *df* | *Deviance* | *p – value* |
| --- | --- | --- | --- | --- |
| **Het1 \* Het 2** | **-596.60** | **1** | **40.256** | $<< 0.01$ |
| | | **3** | **42.724** | $<< 0.01$ |
| Het 1 + Het 2 | -616.73 |  |  |  |
| Null Model | -617.96 |  |  |  |

The strong negative relationship between the abundance of the dominant ant species and the richness of other species (Figure 2) could have two explanations. First, the presence of the dominant species could have precluded the presence of other species, and the more conspicuous this presence was, the more the site was avoided by other species. In the second scenario, the numerically dominant species may have avoided sites where there was a high number of other species. However, we have reasons to think the first scenario is more plausible. There is evidence that numerical dominance is associated with behavioural dominance (Hölldobler & Wilson, 1990) and even that aggressiveness can increase in ants that can perceive their numerical dominance (Tanner, 2006). However, there is no evidence that an ant species is capable of detecting the number of different species that forage in a site and that it might avoid sites where this number is too high. Nevertheless, we think that the most important aspect of our results is that they might shed light on why heterogeneity hypotheses are not completely supported by empirical data. We think that the system in which we executed our study reveals important aspects of this question in relation to scale and indirect dominance effects.

The heterogeneity hypothesis was first proposed and extensively discussed for large-scale conditions (Blackwell, 2007; MacArthur & Wilson, 1963; Tews et al, 2004). At larger scales, the effect of heterogeneity is perceived mostly at the population level, and therefore, processes related to populations can be addressed to understand the partial role of heterogeneity in determining species distribution patterns. For example, Tamme et al (2010) suggested that negative relationships between heterogeneity and richness might be due to fragmentation effects. They argued that as heterogeneity increases on the landscape scale, fragmentation might also increase; therefore, richness might decrease. The loss of species due to fragmentation effects is related to a loss of habitat area for a population, which reduces population size and, in turn, increases the chance of extinction (Fahrig, 2003; Saunders et al, 1991). This mechanism is not reasonable on scales in which heterogeneity is determined by small structures and not by patches of environments that may shelter entire populations.

We think that the patterns observed in our results reveal mechanisms that occur at small scales and that might explain non-positive relationships between heterogeneity and species richness. For example, on small scales, it is possible to observe what Andersen (1992) called “momentary diversity”. This diversity reflects the behaviour of individuals over a short span of time. Andersen explained that the distribution of species changes on local scales in response to the presence of dominant species. Because the abundance of the dominant species changes very quickly, so does the distribution of the species. If the abundance of the dominant ants varies independently from heterogeneity, a survey on the system might show no relationship between heterogeneity and species richness.

In our study, each trap unit was slightly larger than the conventional foraging range of a single nest (10 m × 10 m) (Carroll & Janzen, 1973; Gordon, 1995; Harrison & Gentry, 1981). Heterogeneity on this scale could have effects on nesting and foraging behaviour, but it seems unlikely that an increase in heterogeneity would cause any fragmentation effects. This indicates that the “momentary diversity” effect might be the process behind the pattern we observed.

The effect of heterogeneity mediated by dominance effects also could reveal new explanatory mechanisms about negative relationships between richness and heterogeneity. In the present study, both measures of heterogeneity could either have a positive or negative effect on abundance of the dominant species distribution depending on the value of the other measure (see Figure 3) (Fitzmaurice, 2000). However, there is a negative relationship between Heterogeneity 1 and abundance of dominant species, for mean values of Heterogeneity 2. And there is a positive relationship between Heterogeneity 2 and richness, for mean values of Heterogeneity 1. There is support in the literature to both patterns. Ants have high demand of carbohydrates, protein and heat (Hölldobler &Wilson, 1990). Environments with high values of Heterogeneity 2 might have variability of resource enough to provide all necessities of the dominant population. Therefore, foraging in these environments could be more energy-efficient than foraging in environments with low values of Heterogeneity 2. On the other hand, it has been suggested that structurally dense sites (high values of Heterogeneity 1) are not preferable to some ant species because these habitats might be more energy-consuming to navigate than more homogeneous habitats (see the size-grain hypothesis of Kaspari & Weiser, 1999).

**Figure 3.**
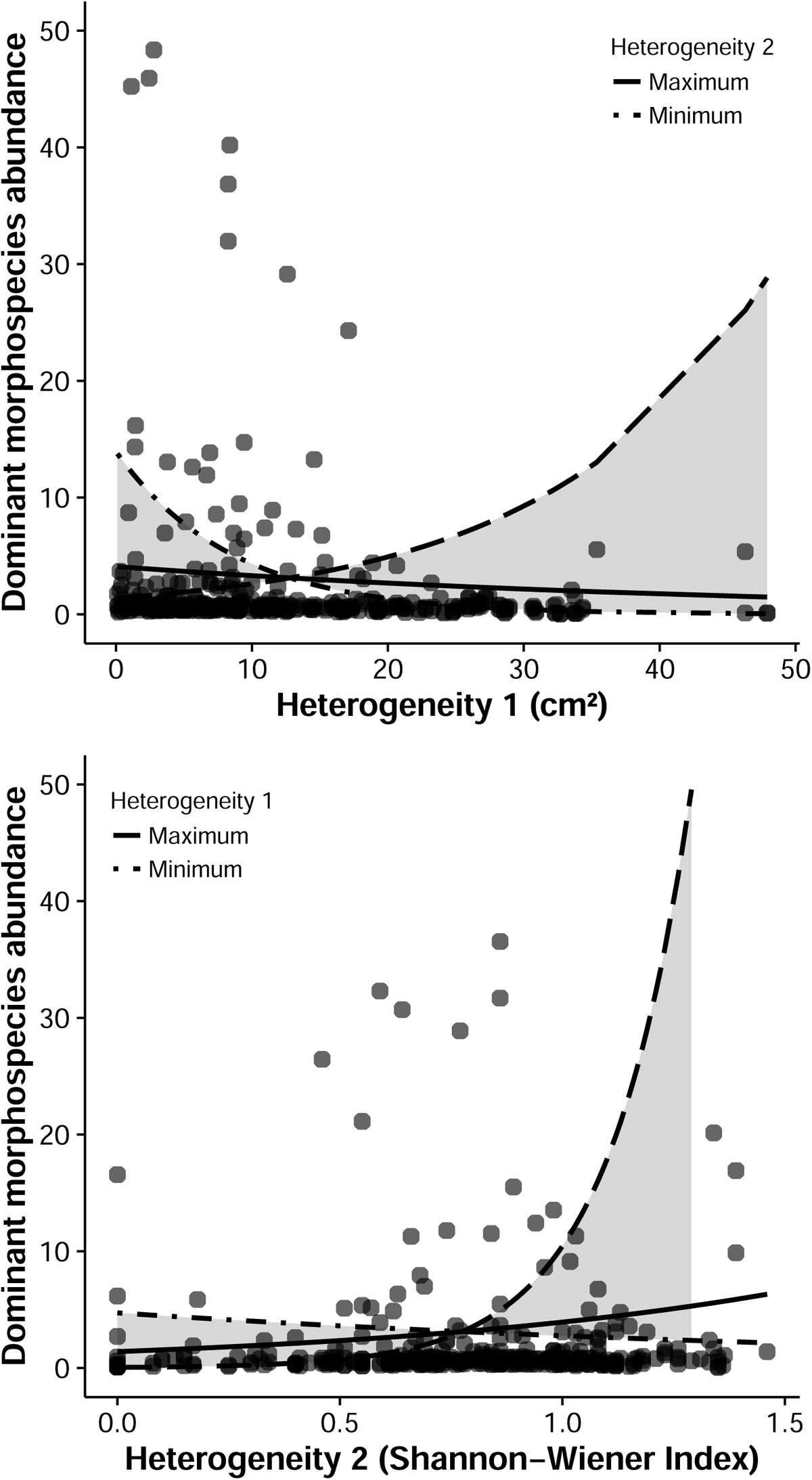
Poisson model fit to the abundance data of the dominant species in relation to Heterogeneity 1 and its interaction with Heterogeneity 2 (top) and vice-versa. Both graphs show the relationship between the heterogeneity measure in the ‘x’ axis and the abundance of the dominant species for the mean value of the other measure (solid line), maximum value observed (long-dash line), and minimum value observed (dash-dot line). The shaded area shows this variation in a continuum from minimum to maximum value.

The relationships between heterogeneity and abundance of the dominant species could revel some aspects of the relationship between heterogeneity and richness in small scales. In our study the abundance of the dominant species was negatively associated with species richness (see Figure 2). The abundance of dominant species was also associates with structurally dense environments (see Figure 3 top). If the dominant species frequently forages in less structurally dense environments, a higher number of species in more structurally dense sites (higher values of Heterogeneity 1) could be explained by the intermediation of the dominance effect. The submissive species are being “pushed” into more structurally dense sites by the dominant species. The same logic can be applied to situations in which the dominant species is positively associated with heterogeneity (see Figure 3 bottom). Dominant species might be pushing submissive species to sites with less diversity of structures. This could explain how species richness is negatively associated with heterogeneity at smaller scales.

Another possibility is that the dominance structure in some systems might be more complex than we conceive. To generate our dominance model, we considered the simplest dominance system, which has only dominant and submissive ants. Dominance relationships in ants can have other elements, including sub-dominant ants. Arnan et al (2011) showed that in some cases, the effect of the presence of a dominant species on the richness of other species is positive. They argued that a high abundance of a dominant species precludes the occurrence of sub-dominant species, which allows for much more submissive species to occur. In these cases, if the dominant ants forage in the more homogeneous environment, we might expect to find a negative relationship between heterogeneity and ant species richness.

Dominance is a very important characteristic in determining ant species distribution patterns (Hölldobler &Wilson, 1990). In other taxa for which dominance relationships are not so important, the pattern we observed might not be so clear. In our study, the strong effect of dominance when compared with that of heterogeneity may not be a particularity of the dominance structure of the taxon studied but instead a particularity of the environment where the study was carried out. In the semi-desert environment of these dunes, resources are very limited, which should intensify the effects of competition in ants (Brown et al, 1979), including those concerning dominant aggressive species (Gordon, 1991). However, in relatively homogeneous environments, such as sand dunes, small changes in the structural configuration of the environment can cause large changes in microhabitat variables (Blackwell, 2007; Rosenzweig & Winakur, 1969), which could also lead to strong effects of habitat heterogeneity. This supports the idea that the dominance effect was strong in the current study because of the taxonomic group and not because of the environmental characteristics.

Other result worth noting is the interaction among heterogeneity measures. Although this interaction did not affect richness distribution (Table 1), it affected abundance of dominant species (Table 2). It is interesting that the model in which the heterogeneity measures did not interact was not significantly different from the null model. This results suggests that the effect of the amount of structures on the population is conditioned by the diversity of structures, and vice-versa (Fitzmaurice, 2000). This is another indicative that the method one measure heterogeneity can drastically affect the results of the study (Tews et al, 2004; Travassos-De-Britto & Rocha, 2013). Even though these two features of the habitat interact they have different mechanisms affecting biodiversity. We suggest that amount of structures and diversity be considered separated components of what we may call heterogeneity. As different components they affect biodiversity by different mechanisms. Authors dealing with the habitat heterogeneity hypothesis should take this in consideration.

We conclude that the structure of interspecific relationships might be more important than variables related to structural heterogeneity in determining species distribution patterns. In some situations, interspecific interactions might even be the most important factors, completely masking the effects of heterogeneity. We emphasize that these types of relationships must be taken into consideration when trying to understand the effects of environmental conditions on species distribution. Furthermore, we suggest that future studies incorporate the hypothesis of the effects of heterogeneity mediated by the effects of dominance as elaborated in this discussion.

## ACKNOWLEDGEMENTS

We thank the Laboratório de Vertebrados Terrestres research group for their help during our fieldwork. We are indebted to the Laboratório de Ecologia Teórica research group for their logistical and intellectual support in the production of the manuscript. B. Travassos-de-Britto and P. L. B. Rocha were supported by scholarships from the Conselho Nacional de Desenvolvimento Científico e Tecnológico (CNPq) and Coordenação de Aperfeiçoamento de Pessoal de Nível Superior (CAPES) during this project. The Pró-Reitoria de Pós-Graduação UFBA (PROPG-UFBA) provided financial support for the costs of the English revision service.

